# The left dorsal stream causally mediates the tone labeling in absolute pitch

**DOI:** 10.1101/2020.07.16.206284

**Authors:** Lars Rogenmoser, Andra Arnicane, Lutz Jäncke, Stefan Elmer

## Abstract

**Background:** Absolute pitch (AP) refers to the ability of effortlessly identifying given pitches without the reliance on any reference pitch. Correlative evidence suggests that the left posterior dorsolateral prefrontal cortex (DLPFC) is responsible for the process underlying pitch labeling in AP.

**Objective:** Here, we aimed at investigating the causal relationship between the DLPFC and the pitch-labeling process underlying AP.

**Methods:** To address this, we measured sight-reading performance of right-handed AP possessors and matched control musicians (*N* =18 per sample) under cathodal and sham transcranial direct current stimulation of the left DLPFC. The participants were instructed to report visually presenting notations as accurately and fast as possible by playing with their right hand on a piano. The notations were simultaneously presented with distracting auditory stimuli that either matched or mismatched them in different semitone degrees.

**Results:** Unlike the control participants, the AP possessors revealed an interference effect in that they responded slower in mismatching conditions than in the matching one. Under cathodal stimulation, half of the time discrepancies between matching and mismatching conditions vanished; specifically, the ones with small up to moderate deviations.

**Conclusions:** These findings confirm that the pitch-labeling process underlying AP occurs automatically and is largely non-suppressible when triggered by tone exposure. The improvement of the AP possessors’ sight-reading performance in response to the suppression of the left DLPFC using cathodal stimulation confirms a causal relationship between this brain structure and pitch labeling.

## Introduction

Absolute pitch (AP) is the ability to effortlessly identify the chroma of a tone without the aid of any reference pitch [1,2]. This ability is sparsely distributed in the population (<1%) [3] but yet bears phylogenetic and ontogenetic significance [4]. There is considerable scientific consensus on its acquisition; namely that AP emerges from an interplay of a certain genetic predisposition and specific environmental inputs and learning factors (i.e., music engagement, language exposure) that operate within a sensitive period during childhood development [5–12]. In contrast, lesser consensus exists on the exact mechanisms and involved brain structures driving AP. One brain structure frequently reported to contribute to AP is the planum temporale (PT) [13–19], a region that covers the superior temporal plane posterior to the Heschl’s gyrus and is involved in language and higher auditory functions [20]. In AP possessors, the PT is likely responsible for a higher resolution in the pitch perception of categories; in other words, the encoding of tones within narrower long-term stored categories [11,19,21,22]. Neuroimaging studies have found an increased leftward asymmetry of the PT [15–17], possibly underlying this so-called “categorical pitch perception” in AP [13,14]. Another brain structure reported to be related to AP is the left posterior dorsolateral prefrontal cortex (DLPFC). This area drives conditional associative learning and memory [23–28]. In the context of AP, the posterior DLPFC may be responsible for the process underlying the association between categorized pitches and verbal labels or other abstract or sensorimotor codes [29–31]; in other words, the pitch-labeling process. Neuroimaging studies revealed that AP possessors exhibit comparatively thinner cortical thickness there [32] and selectively recruit this particular brain region during mere tone listening [30]. In the same study, musicians without AP recruited this region while performing a pitch-interval-labeling task. However, they additionally recruited the right inferior frontal cortex (IFG) [30], an area involved in working memory retrieval [33,34]. Conversely, AP possessors did not show this additional activation while performing the same task. This lack of IFG involvement suggests that AP underlies an automatic pitch-labeling process, functioning without the use of working memory resources. A series of experiments using electroencephalography (EEG) confirmed this interpretation and provided corroborative evidence for the specificity of the pitch-labeling process underlying AP. In auditory oddball or labeling tasks, AP possessors displayed absent, reduced or accelerated specific electrophysiological responses pertaining to the P3 complex [35–40], reflecting a more efficient and parsimonious tone processing in AP. Further event-related EEG components representing hierarchically higher cognitive processes were found to be specific and linked to the labeling performance of AP possessors [41].

The purpose of the present study was to investigate the causal role of the left DLPFC in the pitch-labeling process underlying AP by applying a cathodal and sham transcranial direct current stimulation (tDCS) protocol. Cathodal tDCS suppresses cortical excitability of the targeted region, directly diminishing its underlying function [42]. Given that the pitch-labeling process in genuine AP occurs automatically and thus is rather non-suppressible [22,40,43–48], a modulation in the pitch-labeling performance as a result of cathodal stimulation of the left DLPFC would reveal its causal impact on AP. During cathodal and sham stimulation, participants with AP and matched control participants without AP were instructed to sight-read, reporting the presenting notations as accurately and fast as possible by playing with their right hand on a piano. Sight-reading, the practice of reading and immediately performing notations on an instrument or by singing, is an activity that musicians with and without AP easily master. Simultaneously, auditory stimuli were presented during the task that either matched with the notations or mismatched with them. However, these tones were irrelevant to the task. This experimental set-up corresponded to a so-called Stroop paradigm [49,50]. The Stroop paradigm measures the interference in performances resulting from conflicting asymmetrical processes, namely between overlearned automatically running and more effortful processes. The classical Stroop experiments revealed the robust finding of worsened color-naming performance when participants were challenged to name depicted colors of color names semantically standing for different colors [49,50]. In the case of the classical Stroop task, the interference is due to the fact that reading is virtually overlearned among literate people, making the execution of the less familiar practice, namely color naming, demanding when simultaneously suppressing the decoding of the target words. Meanwhile, this Stroop principle was extended to capture interference effects across multiple domains [50] including music cognition [51–53] but also to verify the authenticity of conditions such as synesthesia [54–56] and AP [40,43,47], which both are characterized by non-suppressible uncommon additional experiences that are inaccessible to outsiders.

In accordance with the Stroop paradigm [49,50], we expected that only AP possessors reveal an interference in the mismatching trials, resulting from the distraction in performing the actual sight-reading task due to the unique pitch-labeling process automatically triggered by tone exposure. Thus, this interference is expected to be reflected as a performance drop in the sight-reading activity. Furthermore, we expected this drop to diminish when suppressing the left DLPFC using cathodal stimulation. Should the automatic pitch-labeling process be driven by the left DLPFC, then suppressing its cortical excitability is expected to diminish its function, leading to less interference with the sight-reading activity and thus to an improvement in performing this task.

## Material and methods

### Participants

A sample of 36 healthy musicians participated in this study, of which half were AP possessors (13 females) and the other were non-AP (NAP) control participants (12 females). All participants were right-handed, as determined using the Annett Handedness Inventory [57] and the Edinburgh Handedness Inventory [58]. The two samples were comparable regarding age (*t*_34_ = 1.86, *P* = .07, *d* = .62), the distribution of the sexes (*Χ*^2^_1_ = 0.13, *P* = .72), the general cognitive capability (*t*_34_ = .2, *P* = .84, *d* = .10) as measured by a standard German intelligence screening test (“Kurztest für allgemeine Intelligenz”) [59], and regarding musical aptitude (*t*_34_ = .09, *P* = .93, *d* = .03) as evaluated using the Advanced Measures of Music Audition test [60]. Both AP and NAP participants commenced their musical training at a comparable age range (*t*_34_ = 1.90, *P* = .07, *d* = .63) and trained for a comparable number of years (*t*_34_ = 1.09, *P* = .29, *d* = .47). Six participants of the AP sample and 7 of the NAP sample professionally played the piano as their “first” instrument. This distribution (binary categorization of “pianist” versus “non-pianist”) did not differ between the two samples (*Χ^2^*_1_ = .12, *P* = .72). However, all participants were skilled at playing the piano, as this particular instrument was thought during music education as part of their professional degree program. The values on characteristics and musical background are reported in Table 1. All participants gave written informed consent to a protocol that was approved by the Cantonal institutional review board of Zurich.

**Table 1.** Characteristics and data on the musical background of the two samples.

|  | AP | NAP |
| --- | --- | --- |
| Age (years) | 27.83 (10.14) | 33.56 (8.23) |
| Cognitive capability (IQ scores) | 123.33 (11.25) | 122.05 (13.20) |
| Age at commencement of musical practice (years) | 5.36 (1.55) | 6.67 (2.47) |
| Duration of musical training (years) | 22.47 (9.95) | 26.61 (7.3) |
| Advanced Measures of Music Audition test (percentile) | 83.56 (6.99) | 83.33 (8.56) |

Listed are the means with the standard deviations in brackets. All independent-samples *t*-test calculated for each variable revealed values of $P > .05$ . AP: absolute pitch, NAP: non-absolute pitch (each $N = 18$ ).

### Absolute pitch (AP) verification

AP was confirmed using an established pitch-labeling test previously used in multiple studies on AP [39,61–63]. In this test, the participants were instructed to immediately write down the accordant tonal label of corresponding sine tones (A4 tuned at 440 Hz) presented to them. Hundred and eight tones covering 3 octaves from A3 to A5 were presented in a pseudorandomized order. Each particular tone was presented 3 times, same tones were never presented successively, and each tone had a duration of 1 s. The inter-stimulus interval was of 4 s and filled with Brownian noise. Accuracy was evaluated by summing the number of correct responses. However, the participants were not asked to identify the octaves of the presented tones. The AP possessors performed (% correct responses) considerably better (mean correct: 69.5, *SD* = 21.5) than the NAP participants (mean correct: 9.36, *SD* = 6.9; *t*_20.46_ = 11.29, *P* < 3.04 × 10^−10^, *d* = 3.59). The NAP participants did not perform better than chance level (8.3%; *t*_17_ = .65, *P* = .53). The individual scores are depicted in Figure 1.

**Fig. 1.**
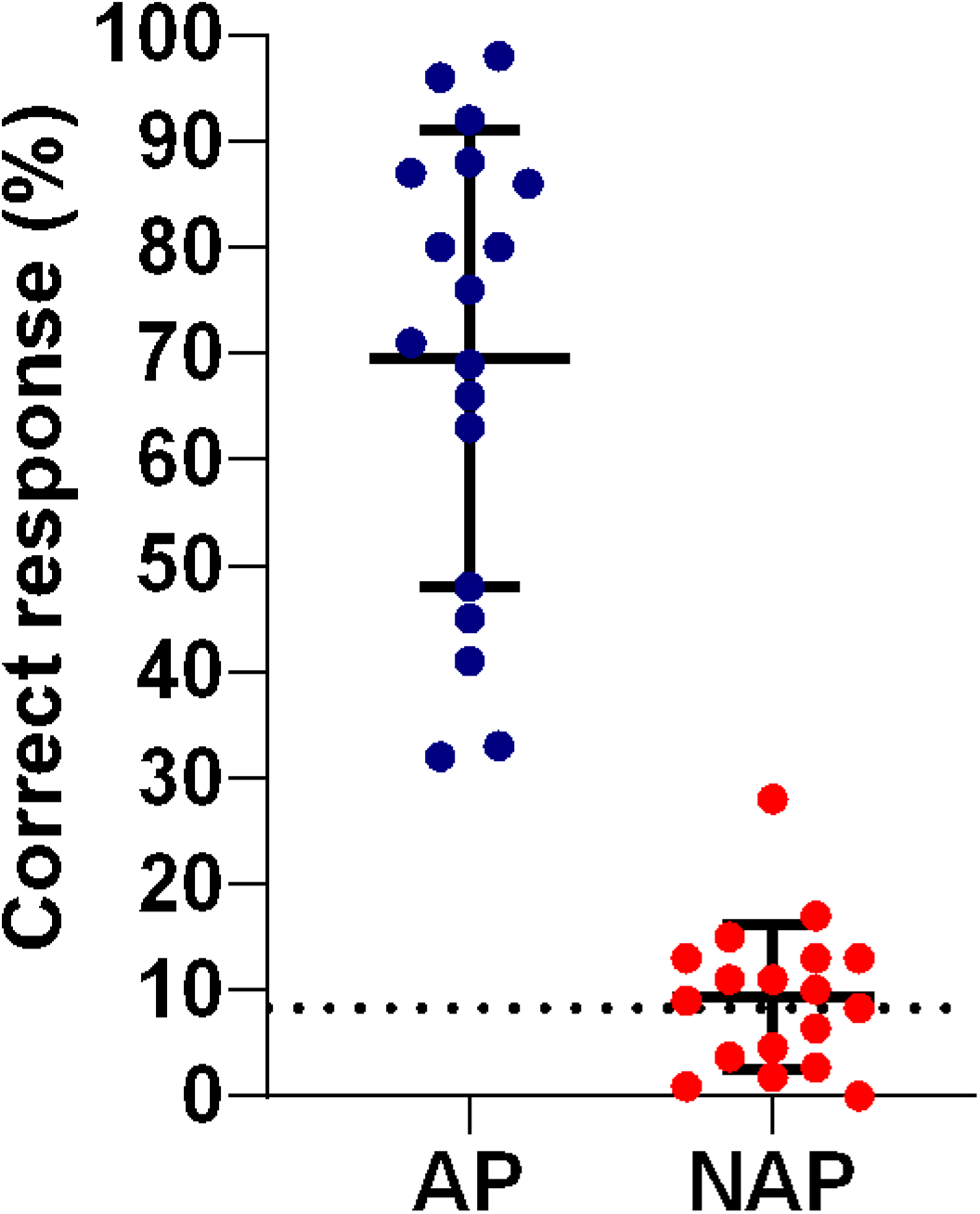
Pitch-labeling performance. Depicted are the individual scores (%) achieved by the participants with absolute pitch (AP, blue circles, *N* = 18) and participants without absolute pitch (NAP, red circles, *N* = 18) from the pitch-labeling test. Plotted are the means with standard deviations. The dotted line represents the baseline at 8.3%.

### Experimental task and stimulus material

The involvement of the left DLPFC in AP ability was investigated by letting both participant samples perform a modified Stroop task under a tDCS protocol. In this task, the participants were exposed to a stream of bimodal musical stimuli, comprising visually presented notations in combination with auditorily presented tones. The participants were instructed to sight-read and report the presenting notations as accurately and fast as possible by playing an electronic piano (Yamaha Electronic Piano, P-60S) with their right hand. During the experiment, the participants positioned their right hand over the piano keys covering the C4 scale in order to be able to respond promptly. The piano did not deliver auditory feedback but recorded the response behavior. The bimodal musical stimuli were randomly presented in different matching conditions: In half of the trials, the tones correctly corresponded to the notations (congruent) whereas in the other half they did not (incongruent). The incongruent trials mismatched in 6 different conditions, deviating between tones and notations in ±1, ±2, and ±3 semitones. Each of the 6 incongruent condition had an occurrence probability of .083. The set of presented notations comprised the C scale of the 12 subsequent notes ranging from C4 to B4. The set of presented auditory stimuli were piano tones corresponding to the particular tones of the C scale (C4 tuned at 262 Hz) including the extension of the 3 successive “deviating” semitones at the scale edges in both directions (A3, B3-flat, B3, and C5, C5-sharp, D5). The tones were professionally recorded with an acoustic piano (University of Iowa Electronic Music Studios, http://theremin.music.uiowa.edu/MISpiano.html) and were trimmed later on, resulting in lengths of 500 ms. In the task, the notations were presented 1000 ms longer than the tones, resulting in a duration of 1500 ms. Afterwards, pink noise followed for a duration of 500 ms. Per trial, responses were allowed and recorded for a duration of < 2 sec after stimulus onset. The inter-trial interval varied randomly between 100-200 ms. The tones were delivered via Sennheiser HD 205 headphones at the sound pressure level of 75 dB, and the notations were shown in the center of a PC monitor mounted on top of the electronic piano. The procedure of one trial is illustrated in Figure 2. Stimulus presentation as well as behavior collection (via Musical Instrument Digital Interface) was controlled by the Presentation software (Neurobehavioral System, Version 18.2).

**Fig. 2.**
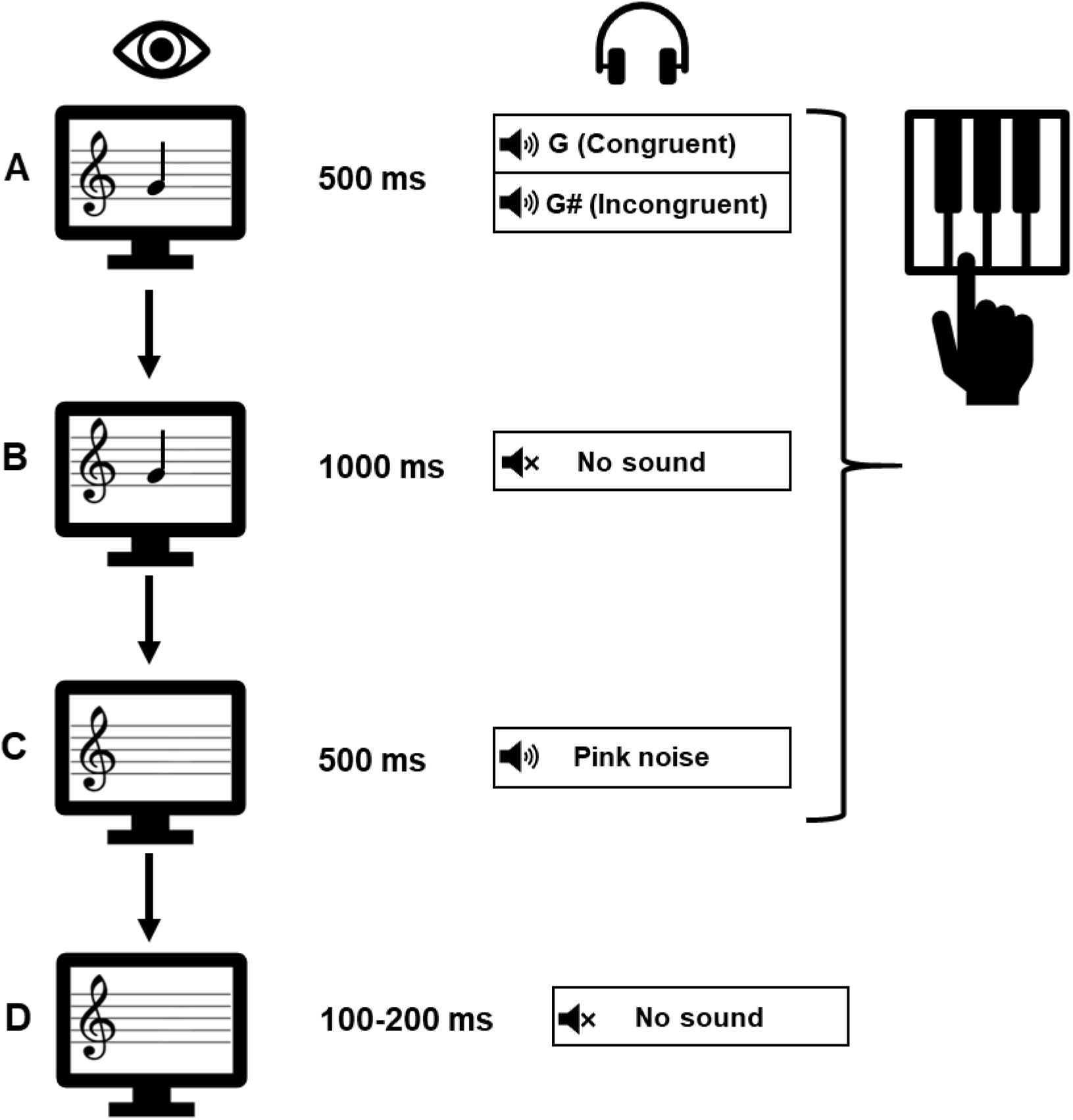
Schematic representation of the task. Each trial began with a bimodal stimulus that lasted for 500 ms (**A**), comprising a notation (e.g., G) presented on a monitor and a piano tone presented via headphones. The piano tone was either congruent (e.g., G) or incongruent (e.g., G#) with the notation. The visual counterpart lasted for an additional 1000 ms (**B**), followed by pink noise for a duration of 500 ms. The next trial followed after a duration jittered between 100 and 200 ms (**D**). The participants were instructed to report the notation as quickly and accurately as possible by sight-reading, specifically by playing with their right hand on a piano. Responses were allowed and recorded for a duration of < 2 s after stimulus onset (**A-C**).

### Transcranial direct current stimulation (tDCS) protocol

After the participants performed a practice block of 20 trials, the tDCS equipment was applied to them. The participants underwent two subsequent experimental blocks; namely one with the tDCS technique turned on, inhibiting the left DLPFC (cathodal stimulation), and one with it turned off (sham stimulation). The order of the blocks was randomized across participants, and the participants were kept unaware of the respective stimulation condition to avoid confounding effects of expectation and order. Each block lasted 10 minutes and consisted of 288 trials, of which half were congruent and the other half incongruent. Regarding the incongruent trials, each combination (12 notations paired with tones mismatching in 6 particular degrees) was presented twice, resulting in 144 trials in total. Regarding the congruent trials, each combination (12 notations paired with matching tones) was presented 12 times, also resulting in 144 trials in total.

A current intensity of 1.5 mA was transferred by a saline soaked pair of surface sponge electrodes and delivered by a battery-driven constant current stimulator (NeuroConn GmnH, Ilmenau, Germany). The sponges were stitched to an electroencephalogram (EEG) cap based on the international “10–20” system to ensure the same placement for all participants. This specific current intensity was chosen based on documentations on the time course of the tDCS after-effect and based on previous tDCS studies investigating the DLPFC [64,65]. For the cathodal stimulation, the current was applied for 9 min including fade-in/out phases of 10 s, respectively. The stimulation and the task were initiated simultaneously. In the sham condition, stimulation was applied for 30 s including fade-in/out phases of 10 s, respectively, so that the subjects experienced the initial tingling associated with onset of stimulation. Between the two blocks, a break of 30 min was held to insure an unaffected sham session [42]. The left DLPFC was targeted by placing the (cathodal) electrode with a surface of 4×8 cm into a sponge on the scalp at the coordinate F3 according to the international “10–20” system. This localization method was successfully conducted in previous studies [65–70], and was confirmed as an appropriate approach by neuronavigational techniques [71]. As reference, an (anodal) electrode with a larger surface of 10×10 cm was placed on the right parietal area, fixating the corners of the sponge at the coordinates Cz, C4, POZ and P8 according to the international “10–20” system. This larger surface size was used to minimize current density over the parietal cortex.

### Data analysis

For each participant, the mean reaction times (RT) and accuracy scores were obtained from each matching and stimulation condition. During the entire study, only 13 missing responses occurred which were omitted. The RT and accuracy values were imported into the SPSS software for statistical analyses. The accuracy scores were not further subjected to inferential statistics due to an obvious ceiling effect (see Table 2). Given that the values of the RTs were largely not normally distributed (see Table 3), as assessed by the Shapiro–Wilk test, non-parametric procedures were performed to determine the effects of interest. For each group (i.e., AP, NAP) and stimulation condition (i.e., sham, cathodal), Friedman tests with “matching” as within-factor (i.e., seven levels: 0, ±1, ±2, ±3 semitones) were run. Significant results were followed up with pairwise comparisons using Wilcoxon signed-rank tests. The Bonferroni procedure was applied to correct for multiple comparisons (corrected α’ < .05/21 = .0024). Effect size measures were calculated; namely the Cohen’s *d* for the *t*-tests, Kendall’s coefficient of concordance (*W*) for the Friedman tests, and the rank-biserial correlation coefficient (*r*) for the Wilcoxon signed-rank tests.

**Table 2.**
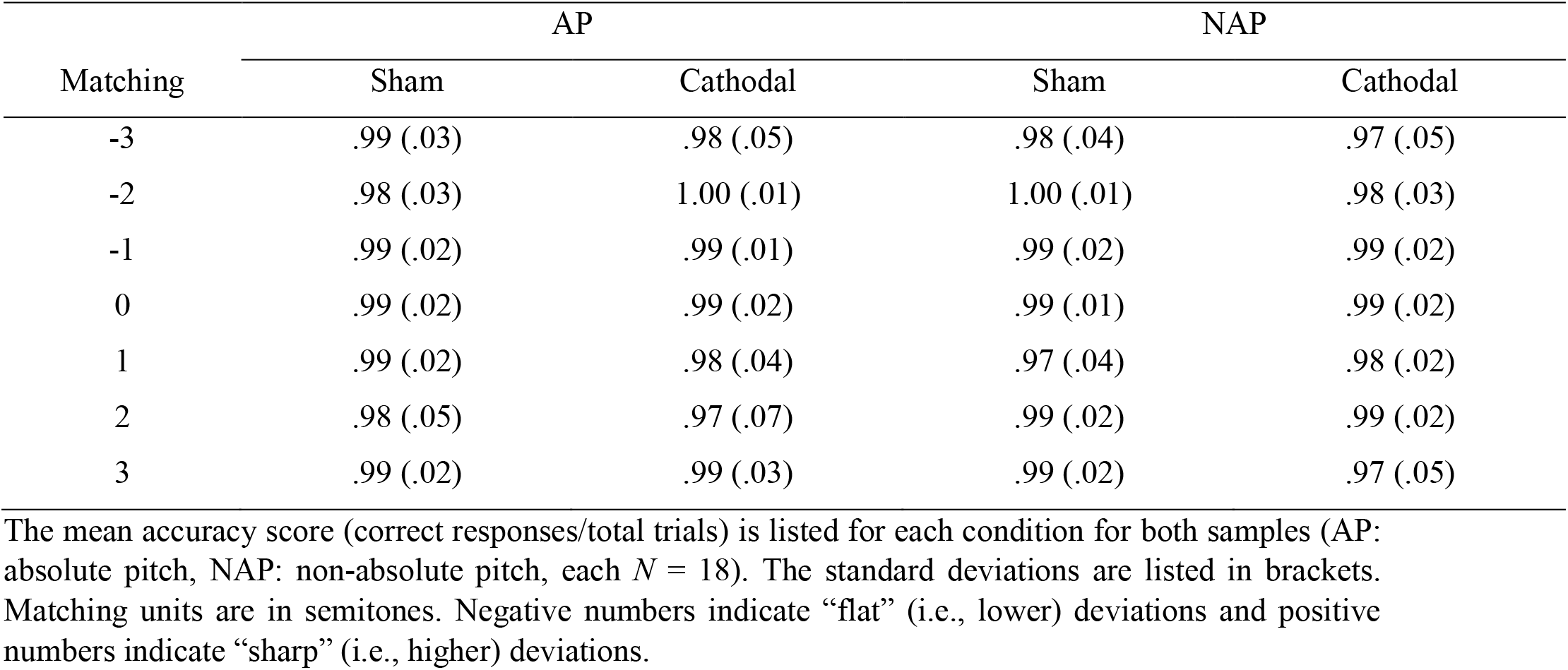
Accuracy scores achieved in the modified Stroop task.

| Matching | AP |  | NAP |  |
| --- | --- | --- | --- | --- |
|  | Sham | Cathodal | Sham | Cathodal |
| -3 | .99 (.03) | .98 (.05) | .98 (.04) | .97 (.05) |
| -2 | .98 (.03) | 1.00 (.01) | 1.00 (.01) | .98 (.03) |
| -1 | .99 (.02) | .99 (.01) | .99 (.02) | .99 (.02) |
| 0 | .99 (.02) | .99 (.02) | .99 (.01) | .99 (.02) |
| 1 | .99 (.02) | .98 (.04) | .97 (.04) | .98 (.02) |
| 2 | .98 (.05) | .97 (.07) | .99 (.02) | .99 (.02) |
| 3 | .99 (.02) | .99 (.03) | .99 (.02) | .97 (.05) |

The mean accuracy score (correct responses/total trials) is listed for each condition for both samples (AP: absolute pitch, NAP: non-absolute pitch, each $N = 18$ ). The standard deviations are listed in brackets. Matching units are in semitones. Negative numbers indicate “flat” (i.e., lower) deviations and positive numbers indicate “sharp” (i.e., higher) deviations.

**Table 3.**
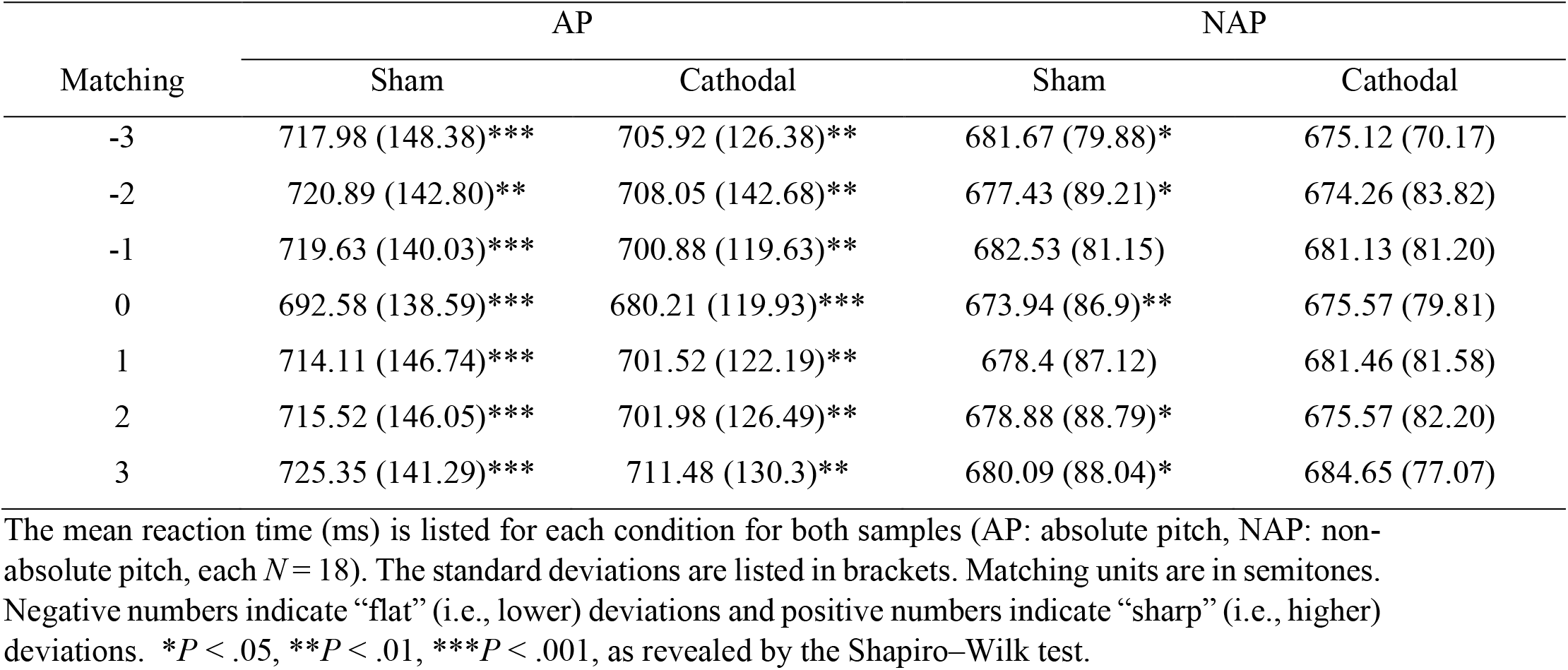
Reaction times achieved in the modified Stroop task.

| Matching | AP |  | NAP |  |
| --- | --- | --- | --- | --- |
|  | Sham | Cathodal | Sham | Cathodal |
| -3 | 717.98 (148.38)*** | 705.92 (126.38)** | 681.67 (79.88)* | 675.12 (70.17) |
| -2 | 720.89 (142.80)** | 708.05 (142.68)** | 677.43 (89.21)* | 674.26 (83.82) |
| -1 | 719.63 (140.03)*** | 700.88 (119.63)** | 682.53 (81.15) | 681.13 (81.20) |
| 0 | 692.58 (138.59)*** | 680.21 (119.93)*** | 673.94 (86.9)** | 675.57 (79.81) |
| 1 | 714.11 (146.74)*** | 701.52 (122.19)** | 678.4 (87.12) | 681.46 (81.58) |
| 2 | 715.52 (146.05)*** | 701.98 (126.49)** | 678.88 (88.79)* | 675.57 (82.20) |
| 3 | 725.35 (141.29)*** | 711.48 (130.3)** | 680.09 (88.04)* | 684.65 (77.07) |

The mean reaction time (ms) is listed for each condition for both samples (AP: absolute pitch, NAP: non-absolute pitch, each $N = 18$ ). The standard deviations are listed in brackets. Matching units are in semitones. Negative numbers indicate “flat” (i.e., lower) deviations and positive numbers indicate “sharp” (i.e., higher) deviations. \* $P < .05$ , \*\* $P < .01$ , \*\*\* $P < .001$ , as revealed by the Shapiro–Wilk test.

## Results

The mean accuracy scores and RTs achieved at each condition by both samples are listed in Table 2 and 3.

Regarding the RTs achieved by the AP possessors, the Friedman test revealed highly significant effects of matching in both stimulation conditions (sham: *Χ^2^*_6_ = 29.64, *P* < .001, *W* = .27; cathodal: *Χ^2^*_6_ = 18.02, *P* = .006, *W* = .17). In the sham condition, post hoc Wilcoxon signed-rank tests revealed significant differences in all possible congruent-incongruent pairs (−3: *z* = −3.72, Bonferroni-adjusted *P* = .004, *r* = −.88; −2: *z* = −3.46, Bonferroni-adjusted *P* = .011, *r* = −.82; −1: *z* = −3.55, Bonferroni-adjusted *P* = .008, *r* = −.84; 1: *z* = −3.16, Bonferroni-adjusted *P* = .033, *r* = −.74; 2: *z* = −3.20, Bonferroni-adjusted *P* = .029, *r* = −.75; 3: *z* = −3.32, Bonferroni-adjusted *P* = .018, *r* = −.78). In the cathodal condition, the AP possessors showed significant differences between the matched and three mismatched conditions (−3: *z* = −3.33, Bonferroni-adjusted *P* = .018, *r* = −.78; 2: *z* = −3.11, Bonferroni-adjusted *P* = .039, *r* = −.73; 3: *z* = −3.29, Bonferroni-adjusted *P* = .021, *r* = −.78). The RTs achieved by the AP possessors are depicted in Figure 3. In the NAP sample, the Friedman test did not reveal any significant effects of matching on the RT (sham: *Χ*^*2*^_6_ = 4.07, *P* = .67, *W* = .04; cathodal: *Χ*^2^_6_ = 8.71, *P* = .19, *W* = .08). The RTs achieved by the NAP participants are depicted in Figure 4.

**Fig.3.**
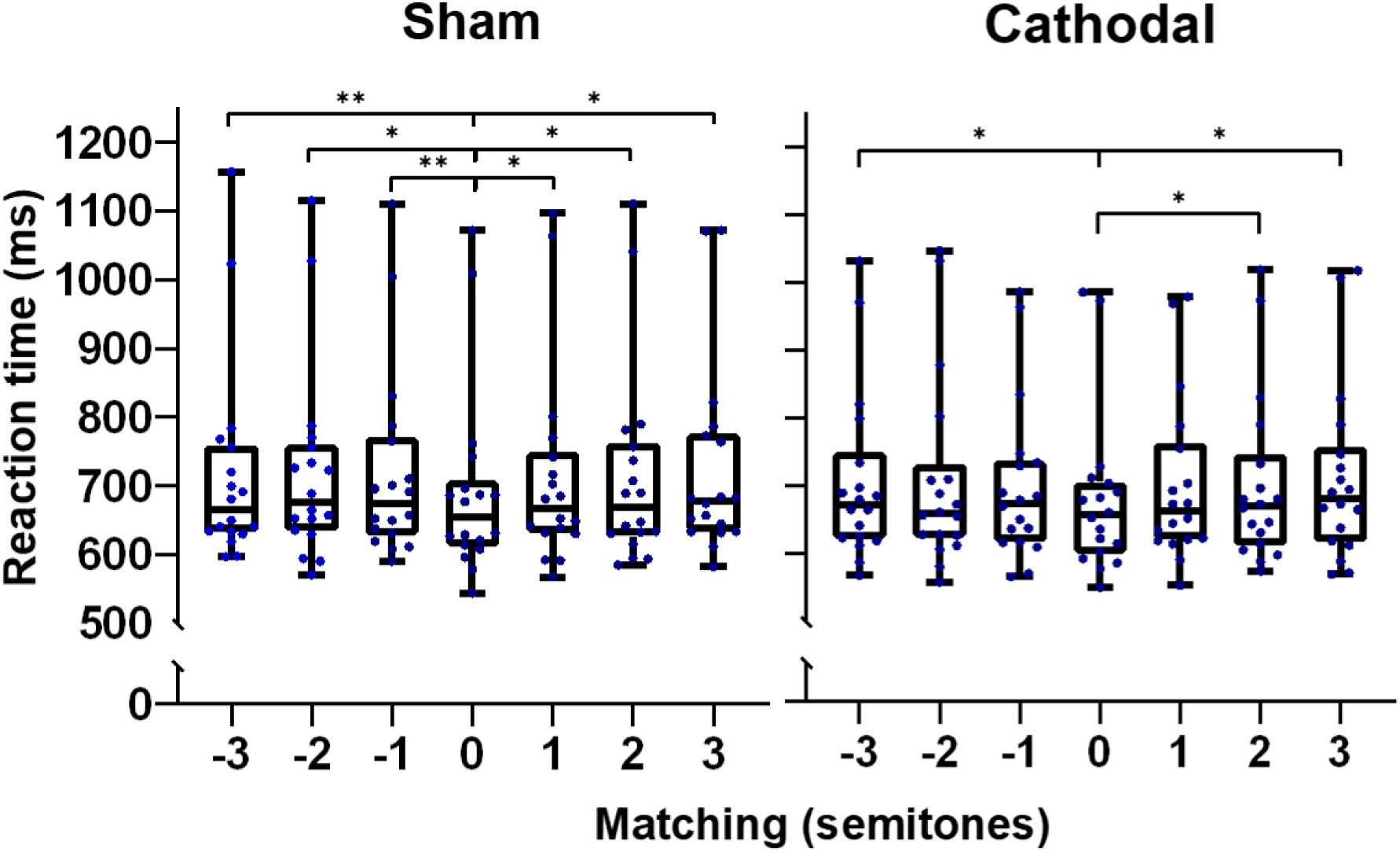
Reaction times achieved by the absolute pitch possessors in the modified Stroop task. Achieved reaction times obtained from each matching and stimulation condition are depicted as boxplots. The blue circles represent the individual values (*N* =18) and the whiskers represent the range. Negative semitones indicate “flat” (i.e., lower) deviations and positive semitones indicate “sharp” (i.e., higher) deviations. Bonferroni-adjusted \*\**P* < .01, \**P* < .05.

**Fig.4.**
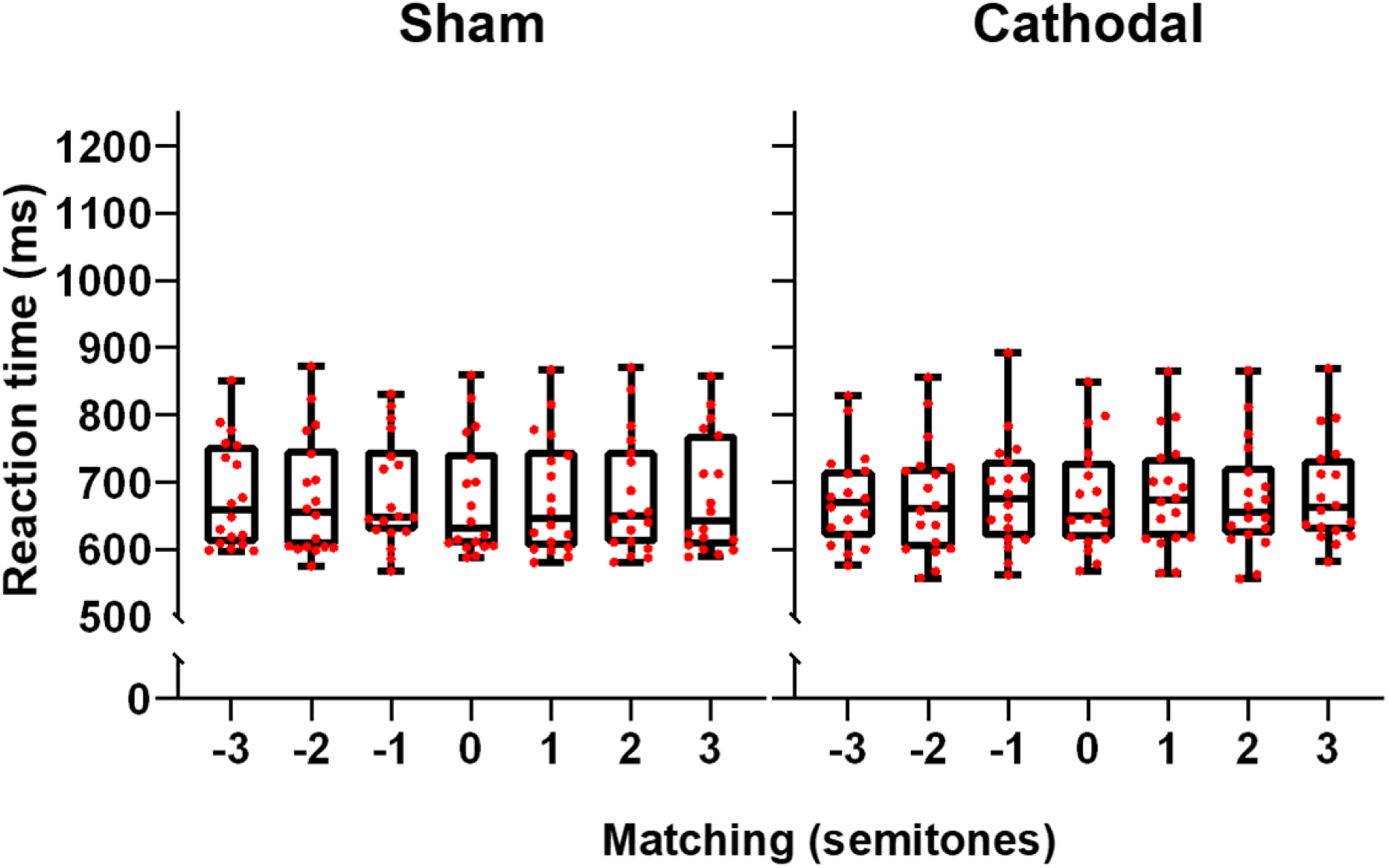
Reaction times achieved by the non-absolute pitch possessors in the modified Stroop task. Achieved reaction times obtained from each matching and stimulation condition are depicted as boxplots. The red circles represent the individual values (*N* =18) and the whiskers represent the range. Negative semitones indicate “flat” (i.e., lower) deviations and positive semitones indicate “sharp” (i.e., higher) deviations.

## Discussion

In this study, we investigated the causal role of the left DLPFC in the pitch-labeling process underlying AP by using a customized Stroop task in the context of a cathodal-sham tDCS protocol. Whereas the previous studies on AP using Stroop tasks recorded the participants’ responses vocally [40,43,47] or per button click [72], our experimental set-up allowed a more natural behavior, namely the responding by performing on an actual piano, constituting a highly familiar situation incorporated in a musician’s everyday life.

### Stroop and tDCS findings

The nearly perfect accuracy scores achieved in the modified Stroop task by both possessors and non-possessors of AP ensure that the notations have properly been internalized and that the sight-reading activity has conscientiously been executed. This compliance appears nontrivial due to the large range and variation of “stimulus-response” commands provided during the task. Whereas the previous Stroop studies on AP included only a handful of stimuli with two matching conditions (i.e., congruent and incongruent) [22,40,43,47,72], our task explored an entire scale (i.e., 12 notations and 12 particular piano key responses) with matching and mismatching conditions, systematically covering 3 double-sided levels of deviation (±1, ±2, ±3 semitones). Despite the participants’ high performance, an interference was still clearly detectable under both stimulation conditions in the AP possessors’ RT. This interference was reflected in that the AP possessors responded slower to notations with mismatching tones than to notations with matching tones. By contrast, the control participants showed no variation in the RT as a function of the matching condition. In line with previous Stroop and other interference studies on AP [22,40,43,45–48,73], these results confirm that the pitch-labeling process is largely non-suppressible when triggered by tone exposure, interfering with the conflicting task of sight-reading. The fact that our well-matched control musician sample without AP did not show this interference pattern further suggests that automaticity in the context of pitch labeling is unique for AP.

The modulation of the task performance as a function of the tDCS stimulation of the left DLPFC indicated, at least to some extent, a causal relationship between this specific brain region and the pitch-labeling process. Under the sham condition, the AP possessors responded systematically slower in all incongruent conditions compared to their responses in the congruent condition. Under the cathodal condition, half of the “incongruent-congruent” differences, specifically the ones with small up to moderate deviations (i.e., −2, ±1 semitones), vanished. In other words, the suppression of cortical excitability of the left DLPFC led to a better sight-reading performance in the AP possessors, suggesting less interference and thus that at least some of the irrelevant tones became more suppressible for them. However, the differences involving larger deviations (i.e., +2, ±3 semitones) were unaffected by the cathodal stimulation. Worth mentioning is that the deviations in the “sharp” direction appeared to be more robust against the cathodal stimulation. We may speculate whether AP underlies a certain asymmetrical proneness. Consistent with this presumption are findings showing that some aging AP possessors undergo a distortion in pitch perception that are mostly biased towards the “sharp” direction [74,75].

Given that we did not perform dose-effect manipulations, our outcome does not allow a dissolution on the factors hindering these “incongruent-congruent” differences in RT involving larger deviations from vanishing under the cathodal stimulation. These unaffected conditions may be a result of our tDCS settings or/and be due to an insufficient contribution of the left DLPFC on the pitch-labeling process. The latter implies that other brain structures may likely be involved in this process as well or may perhaps even be indispensable in this regard. Correlative evidence strongly suggests that the PT is another crucial brain structure for AP [15–17,22,62,76,77]. The PT is anatomically connected via the arcuate fasciculus with the DLPFC [78] and, in AP possessors, the left PT functionally interacts with the left DLPFC already at rest [62]. However, the PT was rather reported to specifically be responsible for early AP-related encoding processes such as “categorical pitch perception” [11,19,21,22]. But, its causal contribution to AP as a whole or to pitch-labeling as a subprocess has not been established yet. A few lesion cases of AP possessors have been documented, but in its entirety so far have been inconclusive in this regard. A few patients were able to retain their AP ability after undergoing left or right temporal lobectomy [79–82]. Others either lost their AP ability or underwent a severe “sharp”-aligned distortion after left or right hemispheric strokes [83,84]. In order to advance our understanding on the mechanisms and the involved brain structures driving AP, future studies using techniques in the field of neuromodulation should be conducted, particularly undertaking the PT next for a more systematic investigation on its causal impact on AP.

### Pitch labeling: A case for the “dorsal stream” function

Previous studies on AP yielded results that are not readily reconcilable with the so-called dual-stream models on auditory cortical processing. These models propose that auditory cortical processing pathways are organized dually, namely ventrally (i.e., the ventral or “what” stream) and dorsally (i.e., the dorsal or “where” stream) [85,86]. Whereas the ventral stream processes the identification of non-spatial auditory properties, the dorsal processing stream integrates spatial sensorimotor information, ultimately associating auditory properties with spatial codes and motor commands [85–87]. In terms of AP, some research has identified the ventral processing stream within the temporal lobe [88,89], while others have loosely assigned the “dorsal stream” function to the left DLPFC and related them to the pitch-labeling process [30–32,62,78]. In two previous studies from our research group, we provided evidence for a correlation between the pitch-labeling performance and the functional and structural connectivity within the left dorsal pathway in AP possessors [62,78]. Here, we extended these finding providing causal evidence of the left DLPFC on the pitch labeling. On this basis, we argue more strongly that the pitch-labeling process underlies the “dorsal stream” function. Our first argument concerns the dorsal location of the assigned brain structure itself that not only drives the pitch-labeling process in AP, as revealed by the present findings and previous studies [30,32,62,78], but also bears an associative-integrative function in NAP musicians, non-musicians, and even monkeys while learning or performing certain association tasks [23–28,31]. Further in line with the “dorsal stream” function concerns the “where” dimension of pitch labeling. In the human mind, the linkage between space and pitch is profoundly incorporated; apparent not only in our musical notations based on a vertical mapping system but also in our usage of the word “height” to describe both space and pitch. Intercultural research revealed that pitch labels are internally represented in a spatial systematic order and that an availability of space-pitch mapping may even be of prelinguistic nature [90–94]. In some rare cases (e.g., pitch-space synesthesia), this order may even reach explicitness, consisting of particular unique pitch-location pairs [95,96]. Finally, in line with the “dorsal stream” function is the co-activation of motor commands during pitch labeling. There is evidence that AP possessors not only rely on verbal information during pitch labeling but also on sensorimotor codes (e.g., specific vocalization or fingering unambiguously coupled to specific tone responses) [29]. Consistent with this framework and our findings, AP possessors show an interference when vocally imitating mistuned tones [97] and a stronger left hemispheric activation during the processing of auditory feedback for vocal motor control [98].

## Conclusions

By applying a cathodal-sham tDCS protocol, we provided, for the first time, causal evidence that the left DLPFC drives the pitch-labeling process underlying AP. Furthermore, our findings yield from our customized piano-playing Stroop task support automaticity as a unique feature of AP, confirming a unique pitch-processing mode virtually non-reliant on cognitive load. Altogether, these findings substantiate previous functional studies showing that AP possessors selectively recruit the left DLPFC during tone exposure and label tones without relying on working memory resources [30], as discussed in terms of reduced or absent P3 responses and lack of activation in the IFG [30,35–40].

## Acknowledgements

This work was supported by the Swiss National Science Foundation (138668, 163149 to LJ and 158642, 186636 to LR). We thank Dr. Jürg Kühnis for coding the experiment.

## References

[1] Baggaley J. Measurement of absolute pitch. Psychol Music 1974;2:11–7. doi:10.1177/030573567422002.

[2] Takeuchi AH, Hulse SH. Absolute pitch. Psychol Bull 1993;113:345–61. doi:10.1037/0033-2909.113.2.345.

[3] Gregersen PK, Kowalsky E, Kohn N & ME. Absolute pitch: Prevalence, ethnic variation, and estimation of the genetic component. Am J Hum Genet 1999;65:911.

[4] Levitin DJ, Rogers SE. Absolute pitch: Perception, coding, and controversies. Trends Cogn Sci 2005;9:26–33. doi:10.1016/J.TICS.2004.11.007.

[5] Theusch E, Basu A, Gitschier J. Genome-wide study of families with absolute pitch reveals linkage to 8q24.21 and locus heterogeneity. Am J Hum Genet 2009;85:112–9. doi:10.1016/J.AJHG.2009.06.010.

[6] Baharloo S, Service SK, Risch N, Gitschier J, Freimer NB. Familial aggregation of absolute pitch. Am J Hum Genet 2000;67:755–8. doi:10.1086/303057.

[7] Baharloo S, Johnston PA, Service SK, Gitschier J, Freimer NB. Absolute pitch: An approach for identification of genetic and nongenetic components. Am J Hum Genet 1998;62:224–31. doi:10.1086/301704.

[8] Deursch D, Henthron T, Dolson M. Speech patterns heard early in life influence later perception of the Tritone Paradox. Music Percept 2004;21:357–72. doi:10.1525/mp.2004.21.3.357.

[9] Deutsch D, Henthron T, Dolson M. Absolute pitch, speech, and tone language: Some experiments and a proposed framework. Music Percept 2004;21:339–56. doi:10.1525/mp.2004.21.3.339.

[10] Deutsch D, Henthorn T, Marvin E, Xu H. Absolute pitch among American and Chinese conservatory students: Prevalence differences, and evidence for a speech-related critical period. J Acoust Soc Am 2006;119:719. doi:10.1121/1.2151799.

[11] Zatorre RJ. Absolute pitch: A model for understanding the influence of genes and development on neural and cognitive function. Nat Neurosci 2003;6:692–5. doi:10.1038/nn1085.

[12] Russo FA, Windel DL, Cuddy LL. Learning the “Special Note”: Evidence for a critical period for absolute pitch acquisition. Music Percept 2003;21:119–27. doi:10.1525/mp.2003.21.1.119.

[13] Loui P, Li HC, Hohmann A, Schlaug G. Enhanced cortical connectivity in absolute pitch musicians: A model for local hyperconnectivity. J Cogn Neurosci 2010;23:1015–26. doi:10.1162/jocn.2010.21500.

[14] Jäncke L, Langer N, Hänggi J. Diminished whole-brain but enhanced peri-sylvian connectivity in absolute pitch musicians. J Cogn Neurosci 2012;24:1447–61. doi:10.1162/jocn_a_00227.

[15] Wilson SJ, Lusher D, Wan CY, Dudgeon P, Reutens DC. The neurocognitive components of pitch processing: Insights from absolute pitch. Cereb Cortex 2009;19:724–32.

[16] Keenan JP, Thangaraj V, Halpern AR, Schlaug G. Absolute pitch and planum temporale. NeuroImage 2001;14:1402–8. doi:10.1006/nimg.2001.0925.

[17] Schlaug G, Jancke L, Huang Y, Steinmetz H. In vivo evidence of structural brain asymmetry in musicians. Science 1995;267:699 LP – 701. doi:10.1126/science.7839149.

[18] Burkhard A, Hänggi J, Elmer S, Jäncke L. The importance of the fibre tracts connecting the planum temporale in absolute pitch possessors. NeuroImage 2020;211:116590. doi:10.1016/J.NEUROIMAGE.2020.116590.

[19] Schulze K, Gaab N, Schlaug G. Perceiving pitch absolutely: Comparing absolute and relative pitch possessors in a pitch memory task. BMC Neurosci 2009;10:106. doi:10.1186/1471-2202-10-106.

[20] Griffiths TD, Warren JD. The planum temporale as a computational hub. Trends Neurosci 2002;25:348–53. doi:10.1016/S0166-2236(02)02191-4.

[21] Siegel JA. Sensory and verbal coding strategies in subjects with absolute pitch. J Exp Psychol 1974;103:37–44. doi:10.1037/h0036844.

[22] Schulze K, Mueller K, Koelsch S. Auditory stroop and absolute pitch: An fMRI study. Hum Brain Mapp 2013;34:1579–90. doi:10.1002/hbm.22010.

[23] Petrides M. Nonspatial conditional learning impaired in patients with unilateral frontal but not unilateral temporal lobe excisions. Neuropsychologia 1990;28:137–49. doi:https://doi.org/10.1016/0028-3932(90)90096-7.

[24] Petrides M, Alivisatos B, Evans AC, Meyer E. Dissociation of human mid-dorsolateral from posterior dorsolateral frontal cortex in memory processing. Proc Natl Acad Sci 1993;90:873 LP – 877. doi:10.1073/pnas.90.3.873.

[25] Petrides M. Deficits in non-spatial conditional associative learning after periarcuate lesions in the monkey. Behav Brain Res 1985;16:95–101. doi:10.1016/0166-4328(85)90085-3.

[26] Petrides M. Visuo-motor conditional associative learning after frontal and temporal lesions in the human brain. Neuropsychologia 1997;35:989–97. doi:https://doi.org/10.1016/S0028-3932(97)00026-2.

[27] Boettiger CA, D’Esposito M. Frontal networks for learning and executing arbitrary stimulus-response associations. J Neurosci 2005;25:2723–32. doi:10.1523/JNEUROSCI.3697-04.2005.

[28] Lepage M, Brodeur M, Bourgouin P. Prefrontal cortex contribution to associative recognition memory in humans: an event-related functional magnetic resonance imaging study. Neurosci Lett 2003;346:73–6. doi:https://doi.org/10.1016/S0304-3940(03)00578-0.

[29] Zatorre RJ, Beckett C. Multiple coding strategies in the retention of musical tones by possessors of absolute pitch. Mem Cognit 1989;17:582–9. doi:10.3758/BF03197081.

[30] Zatorre RJ, Perry DW, Beckett CA, Westbury CF, Evans AC. Functional anatomy of musical processing in listeners with absolute pitch and relative pitch. Proc Natl Acad Sci 1998;95:3172–7. doi:10.1073/PNAS.95.6.3172.

[31] Bermudez P, Zatorre RJ. Conditional associative memory for musical stimuli in nonmusicians: Implications for absolute pitch. J Neurosci 2005;25:7718 LP – 7723. doi:10.1523/JNEUROSCI.1560-05.2005.

[32] Bermudez P, Lerch JP, Evans AC, Zatorre RJ. Neuroanatomical correlates of musicianship as revealed by cortical thickness and voxel-based morphometry. Cereb Cortex 2009;19:1583–96. doi:10.1093/cercor/bhn196.

[33] Zatorre RJ, Evans AC, Meyer E. Neural mechanisms underlying melodic perception and memory for pitch. J Neurosci 1994;14:1908 LP – 1919. doi:10.1523/JNEUROSCI.14-04-01908.1994.

[34] Henson RNA, Shallice T, Dolan RJ. Right prefrontal cortex and episodic memory retrieval: A functional MRI test of the monitoring hypothesis. Brain 1999;122:1367–81. doi:10.1093/brain/122.7.1367.

[35] Klein M, Coles M, Donchin E. People with absolute pitch process tones without producing a P300. Science 1984;223:1306 LP – 1309. doi:10.1126/science.223.4642.1306.

[36] Crummer GC, Walton JP, Wayman JW, Hantz EC, Frisina RD. Neural processing of musical timbre by musicians, nonmusicians, and musicians possessing absolute pitch. J Acoust Soc Am 1994;95:2720–7. doi:10.1121/1.409840.

[37] Wayman JW, Frisina RD, Walton JP, Hantz EC, Crummer GC. Effects of musical training and absolute pitch ability on event-related activity in response to sine tones. J Acoust Soc Am 1992;91:3527–31. doi:10.1121/1.402841.

[38] Hantz EC, Crummer GC, Wayman JW, Walton JP, Frisina RD. Effects of musical training and absolute pitch on the neural processing of melodic intervals: A P3 event-related potential study. Music Percept An Interdiscip J 1992;10:25 LP – 42. doi:10.2307/40285536.

[39] Rogenmoser L, Elmer S, Jäncke L. Absolute pitch: Evidence for early cognitive facilitation during passive listening as revealed by reduced P3a amplitudes. J Cogn Neurosci 2015;27:623–37.

[40] Itoh K, Suwazono S, Arao H, Miyazaki K, Nakada T. Electrophysiological correlates of absolute pitch and relative pitch. Cereb Cortex 2005;15:760–9. doi:10.1093/cercor/bhh177.

[41] Elmer S, Sollberger S, Meyer M, Jäncke L. An empirical reevaluation of absolute pitch: Behavioral and electrophysiological measurements. J Cogn Neurosci 2013;25:1736–53. doi:10.1162/jocn_a_00410.

[42] Nitsche MA, Paulus W. Excitability changes induced in the human motor cortex by weak transcranial direct current stimulation. J PhysiolGy 2000;527:633–9. doi:10.1111/j.1469-7793.2000.t01-1-00633.x.

[43] Akiva-Kabiri L, Henik A. A unique asymmetrical stroop effect in absolute pitch possessors. Exp Psychol 2012;59:272–8. doi:10.1027/1618-3169/a000153.

[44] Ziv N, Radin S. Absolute and relative pitch: Global versus local processing of chords. Adv Cogn Psychol 2014;10:15–25. doi:10.2478/v10053-008-0152-7.

[45] Miyazaki K, Rakowski A. Recognition of notated melodies by possessors and nonpossessors of absolute pitch. Percept Psychophys 2002;64:1337–45. doi:10.3758/BF03194776.

[46] Miyazaki K. Absolute pitch as an inability: Identification of musical intervals in a tonal context. Music Percept 1993;11:55 LP – 71. doi:10.2307/40285599.

[47] Miyazaki, K. Interaction in musical-pitch naming and syllable naming: An experiment on a Stroop-like effect in hearing. In: Nakada T., editor. Integrated Human Brain Science: Theory, Method, Application (Music), Amsterdam: Elsevier; 2000, p. 412–23.

[48] Hsieh IH, Saberi K. Language-selective interference with long-term memory for musical pitch. Acta Acust United with Acust 2008;94:588–93. doi:10.3813/AAA.918068.

[49] Stroop JR. Studies of interference in serial verbal reactions. J Exp Psychol 1935;18:643–62. doi:10.1037/h0054651.

[50] MacLeod CM. Half a century of reseach on the stroop effect: An integrative review. Psychological Bulletin 1991;109:163–203. doi:10.1037/0033-2909.109.2.163.

[51] Zakay D, Glicksohn J. Stimulus congruity and S-R compatibility as determinants of interference in a Stroop-like task. Rev Canad Psychol 1985;39:414–23. doi:10.1037/h0080069.

[52] Grégoire L, Perruchet P, Poulin-Charronnat B. The musical stroop effect: Opening a new avenue to research on automatisms. Experimental Psychology 2013. doi:10.1027/1618-3169/a000197.

[53] Stewart L, Walsh V, Frith U. Reading music modifies spatial mapping in pianists. Percept Psychophys 2004;66:183–95. doi:10.3758/BF03194871.

[54] Beeli G, Esslen M, Jäncke L. When coloured sounds taste sweet. Nature 2005;434:38. doi:10.1038/434038a.

[55] Ward J, Huckstep B, Tsakanikos E. Sound-colour synaesthesia: To what extent does it use cross-modal mechanisms common to us all? Cortex 2006;42:264–80. doi:https://doi.org/10.1016/S0010-9452(08)70352-6.

[56] Itoh K, Sakata H, Igarashi H, Nakada T. Automaticity of pitch class-color synesthesia as revealed by a Stroop-like effect. Conscious Cogn 2019;71:86–91. doi:10.1016/j.concog.2019.04.001.

[57] Annett M. A classification of hand preference by association analysis. British Journal of Psychology 1970;61:303–21. doi:10.1111/j.2044-8295.1970.tb01248.x.

[58] Oldfield RC. The assessment and analysis of handedness: The Edinburgh inventory. Neuropsychologia 1971;9:97–113. doi:10.1016/0028-3932(71)90067-4.

[59] Lehrl S. Geistige Leistungsfähigkeit: Theorie und Messung der biologischen Intelligenz mit dem Kurztest KAI 1993.

[60] Gordon E. Manual for the advanced measures of music audiation 1989.

[61] Oechslin MS, Meyer M, Jancke L. Absolute Pitch--Functional Evidence of Speech-Relevant Auditory Acuity. Cerebral Cortex 2010;20:447–55. doi:10.1093/cercor/bhp113.

[62] Elmer S, Rogenmoser L, Kuhnis J, Jäncke L. Bridging the gap between perceptual and cognitive perspectives on absolute pitch. J Neurosci 2015;35:366–71.

[63] Jäncke L, Rogenmoser L, Meyer M, Elmer S. Pre-attentive modulation of brain responses to tones in coloured-hearing synesthetes. BMC Neurosci 2012;13:151.

[64] Nitsche MA, Paulus W. Sustained excitability elevations induced by transcranial DC motor cortex stimulation in humans. Neurology 2001;57:1899–901. doi:10.1212/WNL.57.10.1899.

[65] Elmer S, Burkard M, Renz B, Meyer M, Jancke L. Direct current induced short-term modulation of the left dorsolateral prefrontal cortex while learning auditory presented nouns. Behav Brain Funct 2009;5:29. doi:10.1186/1744-9081-5-29.

[66] Beeli G, Koeneke S, Gasser K, Jancke L. Brain stimulation modulates driving behavior. Behav Brain Funct 2008;4:34. doi:10.1186/1744-9081-4-34.

[67] Fregni F, Boggio PS, Nitsche M, Bermpohl F, Antal A, Feredoes E, et al. Anodal transcranial direct current stimulation of prefrontal cortex enhances working memory. Experimental Brain Research 2005;166:23–30. doi:10.1007/s00221-005-2334-6.

[68] Marshall L, Mölle M, Hallschmid M, Born J. Transcranial direct current stimulation during sleep improves declarative memory. J Neurosci 2004;24:9985–92. doi:10.1523/JNEUROSCI.2725-04.2004.

[69] Marshall L, Mölle M, Siebner HR, Born J. Bifrontal transcranial direct current stimulation slows reaction time in a working memory task. BMC Neurosci 2005;6:23. doi:10.1186/1471-2202-6-23.

[70] Cerruti C, Schlaug G. Anodal transcranial direct current stimulation of the prefrontal cortex enhances complex verbal associative thought. J Cogn Neurosci 2009;21:1980–7.

[71] Herwig U, Satrapi P, Schönfeldt-Lecuona C. Using the international 10-20 EEG system for positioning of transcranial magnetic stimulation. Brain Topogr 2003;16:95–9. doi:10.1023/B:BRAT.0000006333.93597.9d.

[72] Sharma V V, Thaut M, Russo F, Alain C. Absolute pitch and musical expertise modulate neuro-electric and behavioral responses in an auditory stroop paradigm. Front Hum Neurosci 2019;13:932.

[73] Rogenmoser L, Li CH, Jancke L, Schlaug G. Auditory aversion in absolute pitch possessors. BioRxiv 2020:2020.06.13.145029. doi:10.1101/2020.06.13.145029.

[74] Athos EA, Levinson B, Kistler A, Zemansky J, Bostrom A, Freimer N, et al. Dichotomy and perceptual distortions in absolute pitch ability. Proc Natl Acad Sci 2007;104:14795–800. doi:10.1073/pnas.0703868104.

[75] Vernon PE. Absolute pitch: A case study. Br J Psychol 1977;68:485–9. doi:10.1111/j.2044-8295.1977.tb01619.x.

[76] Wengenroth M, Blatow M, Heinecke A, Reinhardt J, Stippich C, Hofmann E, et al. Increased volume and function of right auditory cortex as a marker for absolute pitch. Cereb Cortex 2013;24:1127–37. doi:10.1093/cercor/bhs391.

[77] Ohnishi T, Matsuda H, Asada T, Aruga M, Hirakata M, Nishikawa M, et al. Functional anatomy of musical perception in musicians. Cereb Cortex 2001;11:754–60. doi:10.1093/cercor/11.8.754.

[78] Oechslin MS, Imfeld A, Loenneker T, Meyer M, Jäncke L. The plasticity of the superior longitudinal fasciculus as a function of musical expertise: A diffusion tensor imaging study. Front Hum Neurosci 2010;3. doi:10.3389/neuro.09.076.2009.

[79] Zatorre RJ. Intact absolute pitch ability after left temporal lobectomy. Cortex 1989;25:567–80. doi:10.1016/S0010-9452(89)80018-8.

[80] Suriadi MM, Usui K, Tottori T, Terada K, Fujitani S, Umeoka S, et al. Preservation of absolute pitch after right amygdalohippocampectomy for a pianist with TLE. Epilepsy Behav 2015;42:14–7. doi:10.1016/j.yebeh.2014.10.025.

[81] Schulz R, Horstmann S, Jokeit H, Woermann FG, Ebner A. Epilepsy surgery in professional musicians: Subjective and objective reports of three cases. Epilepsy Behav 2005. doi:10.1016/j.yebeh.2005.07.009.

[82] Usui K, Shinozaki J, Usui N, Terada K, Matsuda K, Kondo A, et al. Retained absolute pitch after selective amygdalohippocampectomy. Epilepsy Behav Reports 2020:100378. doi:https://doi.org/10.1016/j.ebr.2020.100378.

[83] Johannes S, Jöbges ME, Dengler R, Münte TF. Cortical auditory disorders: A case of non-verbal disturbances assessed with event-related brain potentials. Behav Neurol 1998;11:55–73. doi:10.1155/1998/190715.

[84] Wertheim N, Botez MI. Receptive amusia: A clinical analysis. Brain 1961;84:19–30. doi:10.1093/brain/84.1.19.

[85] Hickok G, Poeppel D. The cortical organization of speech processing. Nat Rev Neurosci 2007;8:393–402. doi:10.1038/nrn2113.

[86] Rauschecker JP, Scott SK. Maps and streams in the auditory cortex: Nonhuman primates illuminate human speech processing. Nat Neurosci 2009;12:718–24. doi:10.1038/nn.2331.

[87] Rauschecker JP. Where, When, and How: Are they all sensorimotor? Towards a unified view of the dorsal pathway in vision and audition. Cortex 2018;98:262–8. doi:10.1016/j.cortex.2017.10.020.

[88] Kim SG, Knösche TR. Intracortical myelination in musicians with absolute pitch: Quantitative morphometry using 7-T MRI. Hum Brain Mapp 2016. doi:10.1002/hbm.23254.

[89] Kim SG, Knösche TR. Resting state functional connectivity of the ventral auditory pathway in musicians with absolute pitch. Hum Brain Mapp 2017;38:3899–916. doi:10.1002/hbm.23637.

[90] Bernstein IH, Edelstein BA. Effects of some variations in auditory input upon visual choice reaction time. J Exp Psychol 1971;87:241–7. doi:10.1037/h0030524.

[91] Ariga A, Saito S. Spatial–musical association of response codes without sound. Q J Exp Psychol 2019;72:2288–301. doi:10.1177/1747021819838831.

[92] Jiang Q, Ariga A. The sound-free SMARC effect: The spatial-musical association of response codes using only sound imagery. Psychon Bull Rev 2020. doi:10.3758/s13423-020-01756-1.

[93] Pratt CC. The spatial character of high and low tones. J Exp Psychol 1930;13:278–85. doi:10.1037/h0072651.

[94] Dolscheid S, Hunnius S, Casasanto D, Majid A. Prelinguistic infants are sensitive to space-pitch associations found across cultures. Psychol Sci 2014;25:1256–61. doi:10.1177/0956797614528521.

[95] Akiva-Kabiri L, Linkovski O, Gertner L, Henik A. Musical space synesthesia: Automatic, explicit and conceptual connections between musical stimuli and space. Conscious Cogn 2014;28:17–29. doi:https://doi.org/10.1016/j.concog.2014.06.001.

[96] Linkovski O, Akiva-Kabiri L, Gertner L, Henik A. Is it for real? Evaluating authenticity of musical pitch-space synesthesia. Cogn Process 2012;13:247–51. doi:10.1007/s10339-012-0498-0.

[97] Hutchins S, Hutka S, Moreno S. Symbolic and motor contributions to vocal imitation in absolute pitch. Music Percept 2015;32:254–65. doi:10.1525/MP.2015.32.3.254.

[98] Behroozmand R, Ibrahim N, Korzyukov O, Robin DA, Larson CR. Left-hemisphere activation is associated with enhanced vocal pitch error detection in musicians with absolute pitch. Brain Cogn 2014;84:97–108. doi:10.1016/J.BANDC.2013.11.007.

